# Galectin-1-deficient mice are protected against *Trypanosoma cruzi* infection through altered neutrophil migration and production of reactive oxygen species and nitric oxide

**DOI:** 10.1101/2020.09.11.294199

**Authors:** Thalita Bachelli Riul, Helioswilton Sales de Campos, Djalma de Souza Lima-Junior, Ana Elisa Caleiro Seixas Azzolini, Cristina Ribeiro de Barros Cardoso, Sean R. Stowell, Richard D. Cummings, Yara Maria Lucisano Valim, Marcelo Dias-Baruffi

## Abstract

*Trypanosoma cruzi* is an intracellular parasite that causes Chagas disease that affects millions of people worldwide. Many cellular and molecular aspects of this neglected disease are not fully understood. Prior studies have shown that galectin-1 (Gal-1), a β-galactoside-binding protein that regulates leukocyte recruitment to the inflammatory site, and promotes *T. cruzi* infection, but the mechanism is unclear. Here, we report that C57BL/6 mice lacking Gal-1 (*Lgals1^−/−^*) exhibited lower parasitemia and higher survival rates than their wildtype (WT) counterparts when infected with *T. cruzi* Y strain. Two weeks after infection, *Lgals1^−/−^* mice displayed greater neutrophil accumulation in infection site and heart tissue than WT mice. In *T. cruzi*-infected *Lgals1^−/−^* mice, infiltrated neutrophils produced increased levels of reactive oxygen species (ROS), while macrophages and neutrophils produced increased levels of nitric oxide (NO), which reduced replication and viability of parasites *in vitro* and downregulated IL-1β production. Pharmacological inhibition of NADPH oxidase and NO synthase during early *in vivo* infection reversed the protective effect of Gal-1 deficiency in *Lgals1^−/−^* mice. Together, our findings demonstrate that lacking Gal-1 favors neutrophil migration to the infection site and increases production of ROS and NO, thereby controlling the early steps of *T. cruzi* infection by reducing parasitemia and prolonging survival of infected mice.

## 1. INTRODUCTION

*Trypanosoma cruzi* is an obligate intracellular parasite and the etiological agent of the American trypanosomiasis or Chagas disease. More than 100 years after its identification by Carlos Chagas, this parasitic infection continues to remain a significant social and public health problem in Latin America. Chagas disease is considered a neglected tropical disease by the World Health Organization due to its high prevalence in resource poor regions of the world, including those located in developed countries, as well as to the scarcity of investment for the development of new therapeutic and diagnostic strategies. Recent data estimate that 70 million people live in areas at risk of infection, 57 million people are infected, and 12,000 deaths/year are caused by this anthropozoonosis (Perez-Molina & Molina, 2018). Some factors that contribute to the persistence of *T. cruzi* infection include transmission forms associated with solid organ transplantation, blood transfusion, resurgence of the acute phase in immunosuppressed patients, congenital infection, and high migration of people from endemic to non-endemic regions (Lattes & Lasala, 2014; Martins-Melo, Lima, Ramos, Alencar, & Heukelbach, 2014).

Chagas disease is characterized by an acute phase, which is poorly detected and can appear as a self-limiting febrile illness, followed by an indeterminate/latent phase that can evolve to chronic phase often accompanied by impaired function of cardiac muscle or gastrointestinal tract (Bahia, Diniz, & Mosqueira, 2014; Hotez, 2008; Rassi, Rassi, & Little, 2000). Some features of acute infection, such as parasitemia, extent of initial inflammatory response, and balance of regulatory responses interfere with the likelihood of chronic infection (Borges, Araujo, Cardoso, & Lazo Chica, 2013; Marinho, D’Imperio Lima, Grisotto, & Alvarez, 1999).

Galectin-1 (Gal-1) is a β-galactoside-binding protein with several anti-inflammatory properties. Gal-1 suppresses neutrophil migration to infection sites or a gradient of inflammatory factors, both *in vitro* and *in vivo* (Cooper, Norling, & Perretti, 2008; La et al., 2003; Rabinovich, Sotomayor, Riera, Bianco, & Correa, 2000), only when inflammatory conditions are present (Auvynet et al., 2013). Human recombinant Gal-1 (hrGal-1) inhibits acute inflammation in a mouse model induced by phospholipase A_2_ from bee venom by decreasing neutrophil infiltration (Rabinovich et al., 2000), and suppressing interleukin-1β-induced recruitment of polymorphonuclear cells to mice peritoneal cavity, indicating that this lectin specifically downregulates the initial steps of leukocyte-endothelium interaction (La et al., 2003). In addition, hrGal-1 inhibits neutrophil capture, rolling, and adhesion on activated endothelial monolayers *in vitro* (Cooper et al., 2008). Leukocyte adhesion and emigration are significantly increased in Gal-1-null mice inflamed with IL-1β, as demonstrated by intravital microscopy (Cooper et al., 2008). In contrast, Gal-1 induces neutrophil migration both *in vitro* and *in vivo* in the absence of an inflammatory stimuli (Auvynet et al., 2013). Hence, although the presence of Gal-1 can favor the resolution of adverse inflammatory conditions (Rabinovich et al., 2000; Rodrigues et al., 2016; Seropian et al., 2013), it can be unfavorable in the battle against infection (Davicino et al., 2017). Nevertheless, little is known regarding the role of Gal-1 in parasite infections. A recent study demonstrated that Gal-1 negatively regulates the protective immunity against *T. cruzi*. Compared to Gal-1-null mice, wild-type mice expressing Gal-1 exhibited higher levels of immune inhibitory mediators regulated by tolerogenic dendritic cells, and increased mortality rate and muscle parasitemia (Poncini et al., 2015).

Considering that (i) Gal-1 regulates neutrophil migration to inflammatory sites, (ii) neutrophils play an important role during the early stages of parasitic infection, and (iii) innate immune responses influence the progress of *T. cruzi* infection, in the present study we aimed to examine whether Gal-1 modulates recruitment of peritoneal inflammatory leukocytes and the production of ROS and NO by these cells during the acute phase of the experimental model of *T. cruzi* intraperitoneal infection in C57BL/6 mice.

## 2. RESULTS

To address whether endogenous Gal-1 interferes with the course of acute *T. cruzi* infection, both C57BL/6 WT and *Lgals1^−/−^* mice were infected intraperitoneally with blood trypomastigote forms of *T. cruzi*. The evaluation of health parameters during the acute phase of infection revealed increased blood parasitemia at the parasitemia peak (Figure 1A) and increased mortality rate (Figure 1 B) of WT mice when compared with *Lgals1^−/−^* mice. Interestingly, none of the *Lgals1^−/−^* mice died during the 60-day experimental period (Figure 1B). These data indicate that the presence of endogenous Gal-1 is closely associated with enhanced susceptibility to *T. cruzi* infection. Quantification of the relative amount of parasite DNA in the cardiac tissue evidenced (i) a fourfold increase in parasite DNA levels in WT mice at 14 days of infection, when compared with *Lgals1^−/−^* mice (p<0.001); and (ii) the presence of low and similar levels of parasite DNA in both mice groups at 21 and 28 days of infection (p<0.05) (Figure 1C).

**Figure 1.**
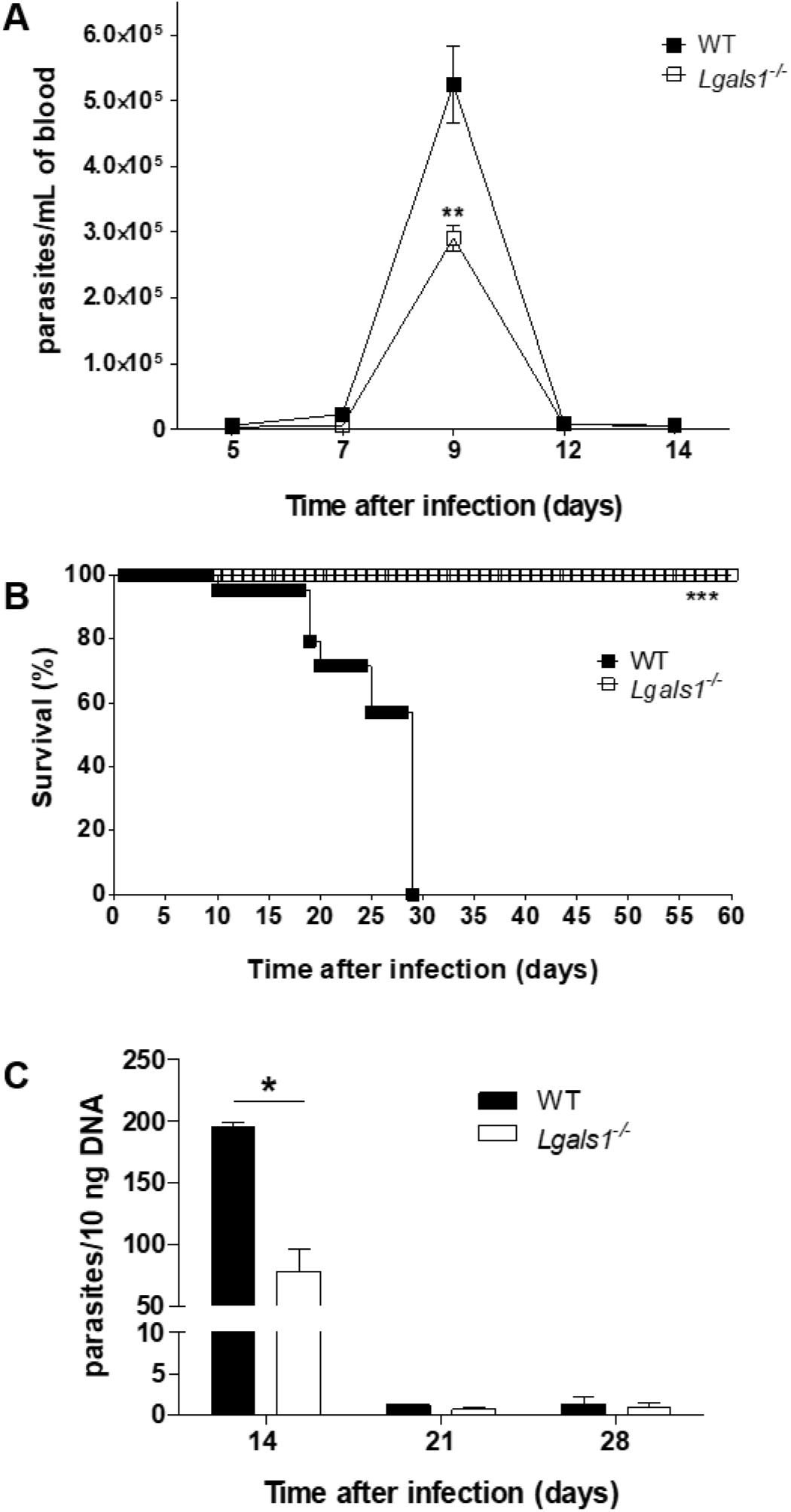
Lack of endogenous Gal-1 increases mice resistance against *T. cruzi* acute infection. (A) Blood parasitemia, (B) Survival rate, and (C) Relative amount of parasites in cardiac tissue (determined by quantitative real-time PCR) from C57BL/6 wild-type (WT) and *Lgals1^−/−^* mice infected intraperitoneally with trypomastigote forms of *T. cruzi* strain Y. Results are expressed as mean ± standard deviation (n = 6). \*\*\**p* < 0.001, \*\**p* < 0.01 and \**p* < 0.05 (unpaired Student *t* test).

As *Lgals1^−/−^* mice exhibited reduced parasitemia in blood and heart tissue during the initial weeks of infection, we characterized the type of immune cells recruited to the peritoneum and heart tissue of WT- and *Lgals1^−/−^-infected* mice, and examined whether they participate in the control of infection at its early steps. At 6, 48, and 72 hours of infection, WT and *Lgals1^−/−^* mice exhibited similar profiles of peritoneal immune cells. There was increased neutrophil infiltration at 6 hours, and increased macrophage infiltration at the three experimental periods. At 24 hours, macrophage infiltration was increased in both mice groups, while neutrophil infiltration was increased only in *Lgals1^−/−^* (Figure 2A). Macrophage and neutrophil infiltration in WT were greater and lower than those detected in *Lgals1^−/−^*, within the first 24 hours of infection. The typical morphology of macrophages, neutrophils, lymphocytes, and mast cells infiltrated into the peritoneal cavity is depicted in Figure 2B.

**Figure 2.**
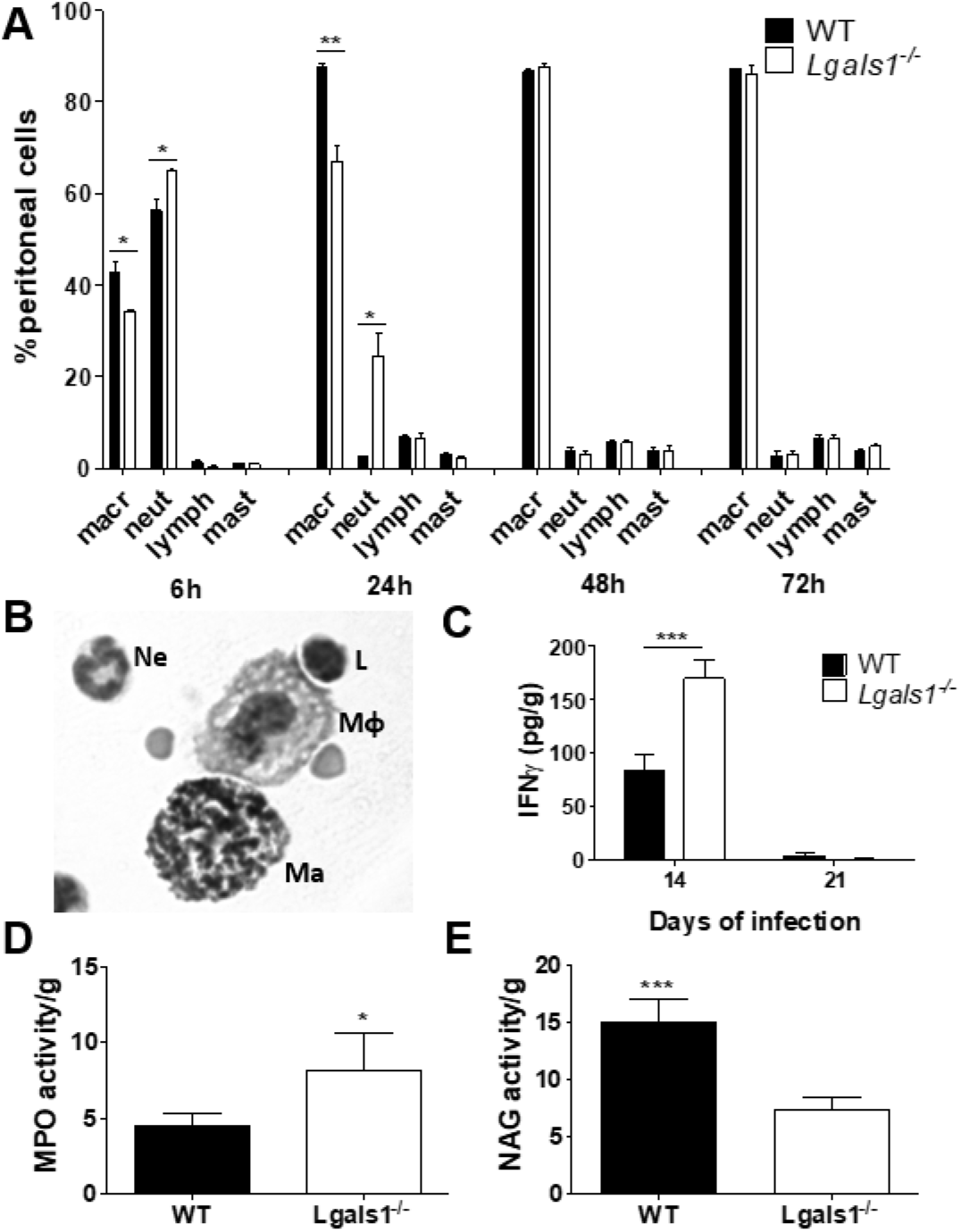
Wild-type and *Lgals1^−/−^* mice acutely infected with *T. cruzi* differ with respect to the cell migration profile into the peritoneal cavity and heart tissue. (A) Relative population of immune cells in the peritoneal cavity determined by differential count, after 6, 24, 48, and 72 hours of infection. (B) Photomicrograph of a typical macrophage (MΦ), neutrophil (Ne), lymphocyte (L), and mast cell (Ma) obtained from peritoneal cavity. 1000x magnification. (C) IFN-γ concentration in heart tissue extract after 14 and 21 days of infection. Determination of (D) myeloperoxidase (MPO) and (E) N-acetyl-β-D-glucosaminidase (NAG) activity to estimate the presence of neutrophils and macrophages in heart tissue after 14 days of infection. Results are expressed as mean ± standard deviation (n = 4-6). \**p* < 0.05; \*\**p* < 0.01; \*\*\**p* < 0.001 (unpaired Student *t* test).

Quantification of interferon-γ (IFNγ) concentration in cardiac tissue extract revealed that its levels were augmented in *Lgals1^−/−^* at 14 days of infection (p<0.001), but it decreased back to the levels detected in WT mice at 21 days of infection (Figure 2C). Determination of the relative number of neutrophils and macrophages infiltrated into cardiac tissue by measuring the activity of MPO and NAG, respectively – specific enzymes from these immune cells – at 14 days of infection demonstrated that *Lgals1^−/−^* had increased MPO activity (Figure 2D), whereas WT had increased NAG activity (Figure 2E). This finding is in line with the enhanced peritoneal neutrophil and macrophage infiltration in *Lgals1^−/−^* and WT, respectively (Figure 2A).

We performed histopathological analysis of cardiac tissue from WT and *Lgals1^−/−^* mice at different time points of *T. cruzi* infection (Figure 3). The integrity of cardiac fibers from both mice groups remained virtually unchanged at 7 days of infection (Figure 3, 7d), when free parasites were still found in the bloodstream (Figure 1A). At 14 days of infection, the foci of inflammatory infiltrate were increased in both mice groups, and amastigote nests seem to accumulate in cardiac tissue from WT mice, but not from *Lgals1^−/−^* mice (Figure 3, 14d). Tissues from WT and *Lgals1^−/−^* mice clearly differed at 21 days of infection: the former still had prominent inflammatory infiltrate, while the latter resembled the tissue from non-infected animals (Figure 3, 21d and NI). Although the photomicrographs did not differentiate the immune cell types present, the aforementioned results on MPO and NAG activity (Figure 2D and 2E, respectively) estimated the relative infiltration of neutrophils and macrophages. In addition, real-time PCR analysis confirmed the highest heart parasitemia in WT mice when compared with *Lgals1^−/−^* mice (Figure 1C).

**Figure 3.**
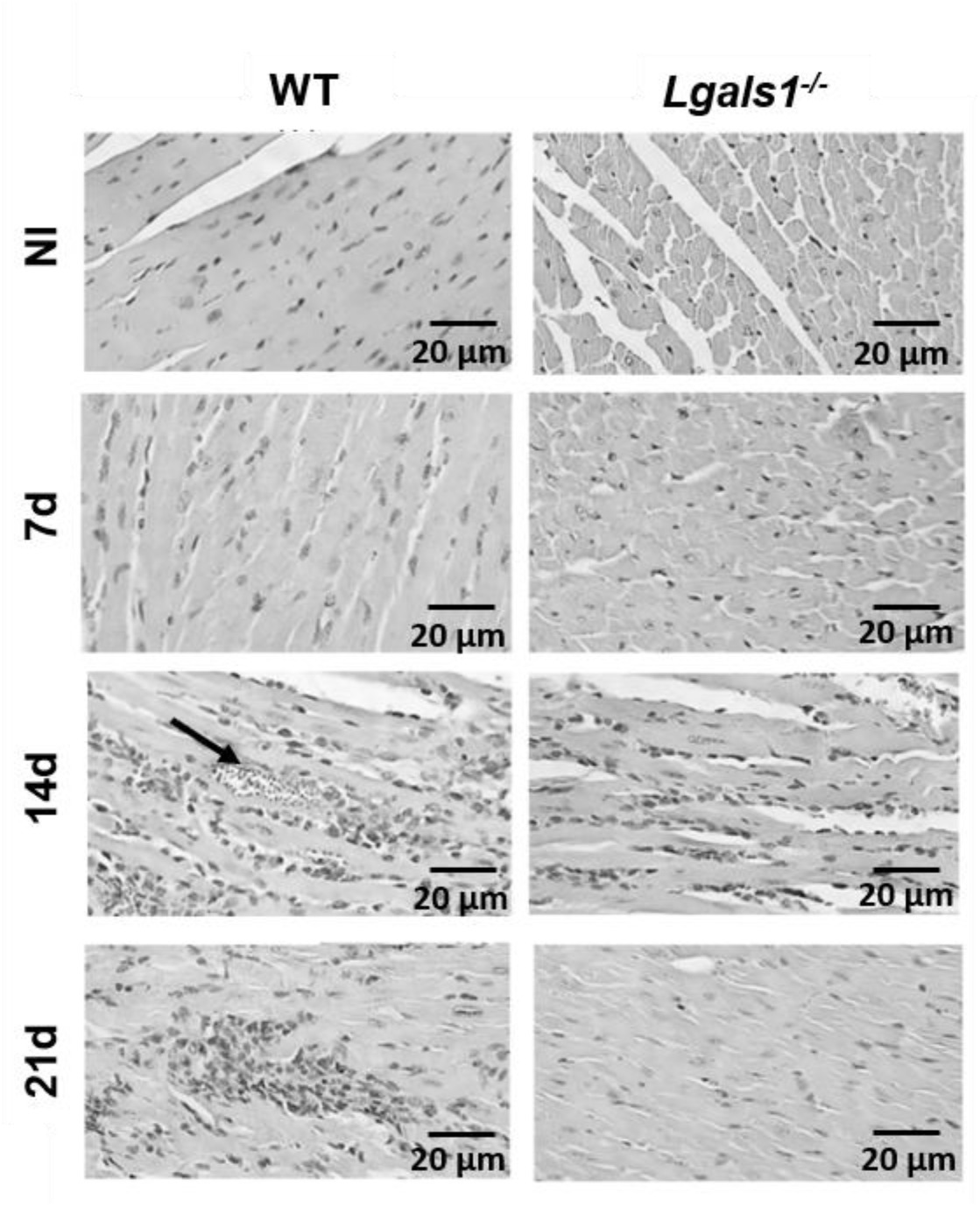
Histology of heart tissue from WT and *Lgals1^−/−^* mice acutely infected with *T. cruzi*. Photomicrographs of representative sections of cardiac tissues from WT and *Lgals1^−/−^* mice collected, processed, and analyzed after 7, 14, and 21 days of infection (7d, 14d, and 21d, respectively). NI: non-infected. The arrow indicates the presence of amastigotes nest.

As neutrophils were the most abundantly recruited cells to the peritoneal cavity and heart tissue of *T cruzi*-infected *Lgals1^−/−^* mice, we analyzed the ROS-producing capacity of these leukocytes, which is an important mechanism for the initial control of infections. Six hours after intraperitoneal infection with *T. cruzi*, leukocytes from the peritoneal cavity of WT and *Lgals1^−/−^* mice were collected, washed, and stimulated with PMA and fMLP to measure ROS production through a chemiluminescence assay. All the cell preparations contained more than 80% of neutrophils. *Lgals1^−/−^* neutrophils produced more ROS than WT mice neutrophils when stimulated with PMA and fMLP (Figure 4A and B, respectively). DPI, an NADPH oxidase inhibitor, suppressed ROS production in fMLP-stimulated neutrophils from *Lgals1^−/−^* mice but not from WT mice (Figure 4B).

**Figure 4.**
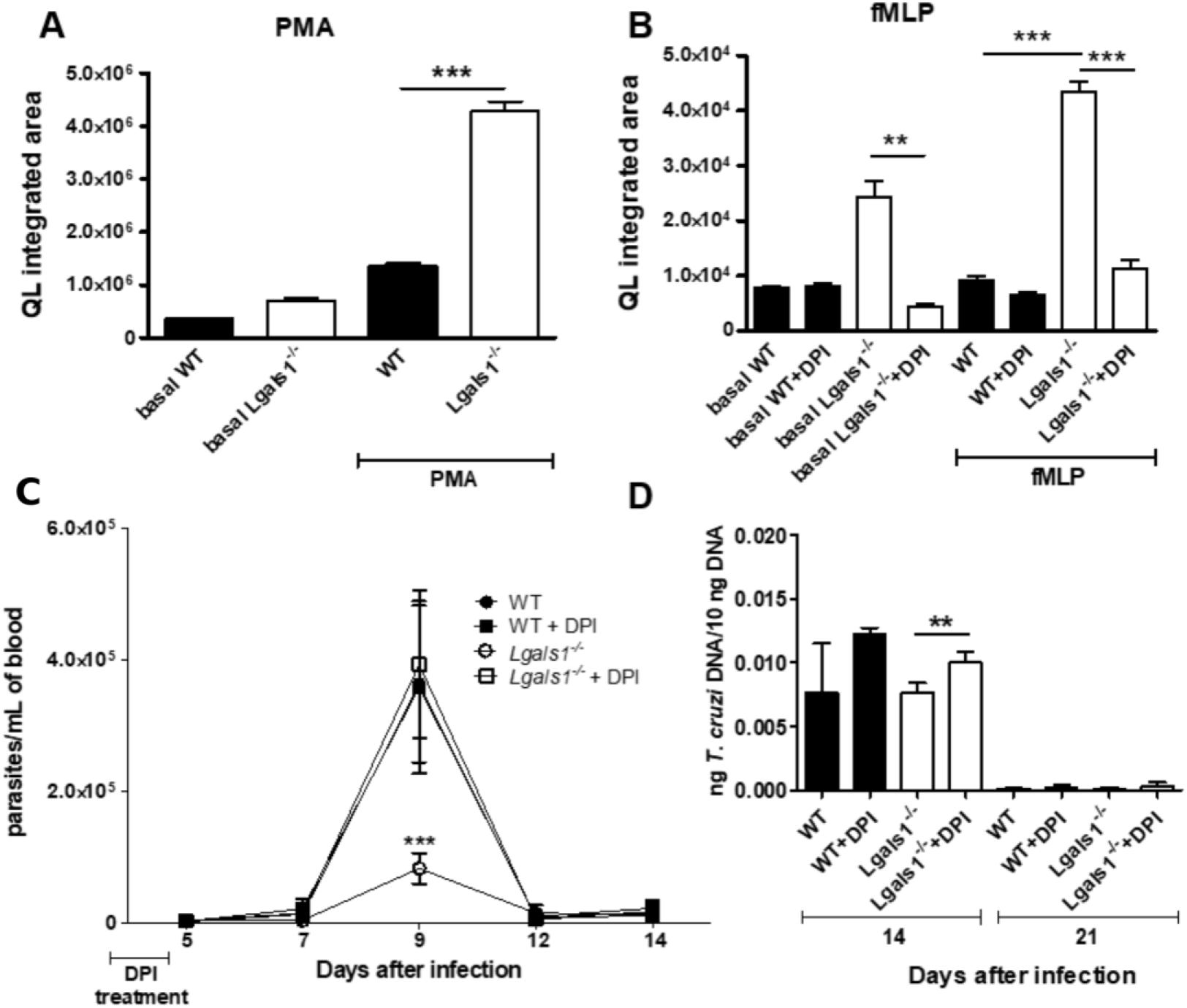
*Lgals1^−/−^* mice acutely infected with *T. cruzi* exhibit stronger peritoneal neutrophil ROS production and control of blood and heart parasitemia, which are reverted by DPI administration. Wild-type (WT) and *Lgals1^−/−^* mice were infected intraperitoneally with *T. cruzi* and, after 6 hours of infection, peritoneal cells (>80% neutrophils) were collected and stimulated with (A) PMA (phorbol 12-myristate 13-acetate) or (B) fMLP (N-formyl-met-leu-phe) to produce ROS, which were detected by the luminol-enhanced chemiluminescence assay. Each bar in (A) and (B) represents a triplicate experiment with pooled cells. Parasitemia in (C) blood and (D) heart tissue was determined after intraperitoneal treatment with DPI during the first five days of infection. *T. cruzi* DNA load in mice heart tissues was determined by quantitative real-time PCR. Each bar in (C) and (D) represents mean ± standard deviation (n = 6 animals). DPI: Diphenyleneiodonium chloride (an NADPH-oxidase inhibitor). \*\**p* < 0.01; \*\*\**p* < 0.001 (unpaired Student’s *t* test; two-way ANOVA combined with Bonferroni *post-hoc* test).

Considering that neutrophils are important cells of the innate immune response for the initial control of infections; *Lgals1^−/−^* mice were resistant to acute *T. cruzi* infection (Figure 1); presented increased neutrophil influx into the peritoneal cavity (Figure 2); and *Lgals1^−/−^* mice neutrophils produced more ROS than WT mice neutrophils (Figure 4A and B). Thus, we examined how inhibition of ROS production in neutrophils early recruited to the peritoneal cavity interfered with the course of acute *T. cruzi* infection. To this end, we administered DPI intraperitoneally to WT and *Lgals1^−/−^* mice during the first five days of infection. At the ninth day of infection, blood parasitemia in DPI-treated *Lgals1^−/−^* mice increased and reached levels similar to those detected in WT mice (Figure 4C). Treatment with DPI did not change blood parasitemia in WT mice. DPI-mediated suppression of ROS production increased parasitemia not only in blood but also in heart tissue of *Lgals1^−/−^* mice after 14 days of infection (Figure 4D). Thus, the rapid influx of highly ROS-producing neutrophils into the infection site correlated with control of the acute *T. cruzi* infection in *Lgals1^−/−^* mice.

As IL-1β is one of the major cytokines that elicits ROS production in macrophages and neutrophils to help control infection (van de Veerdonk, Netea, Dinarello, & Joosten, 2011), we measured its levels in the culture supernatant of these cells from WT and *Lgals1^−/−^* mice cultured *in vitro* with *T. cruzi*. Interestingly, IL-1β levels in cultures of peritoneal neutrophils and BMDM from WT mice were higher than those from *Lgals1^−/−^* mice (Figure 5). Compared with *Lgals1^−/−^* mice macrophages, WT mice macrophages released higher IL-1β levels in response to stimulation with LPS followed or not by infection with *T. cruzi*. (Figure 5B). Hence, IL-1β is unlikely to mediate the enhanced ROS production in *Lgals1^−/−^* mice neutrophils and macrophages.

**Figure 5.**
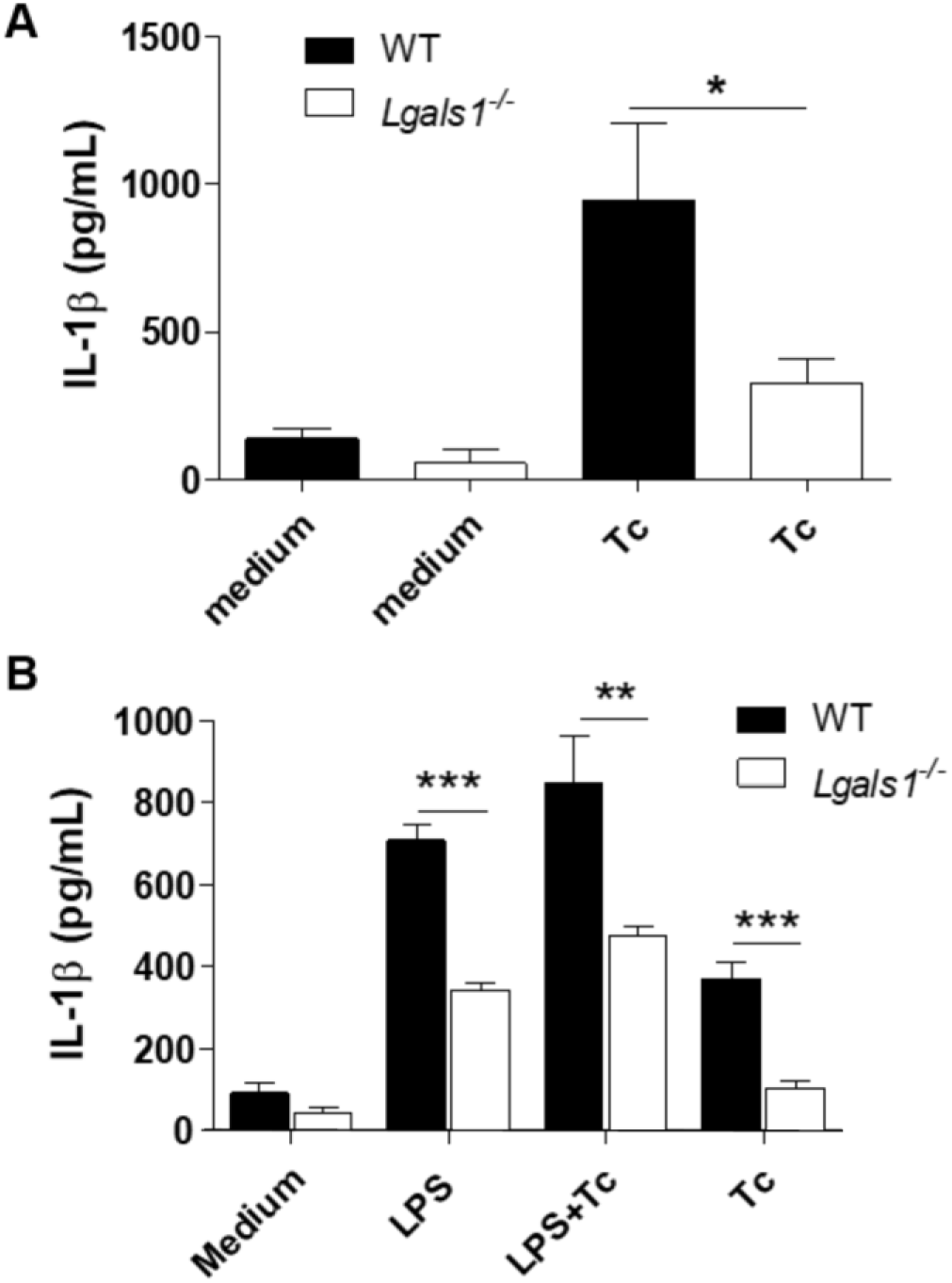
Peritoneal cells and bone marrow-derived macrophages from *T. cruzi*-infected *Lgals1^−/−^* mice produce lower levels of IL-1β. (A) IL-1β levels in the supernatant of peritoneal cells (>80% neutrophils) cultured with *T. cruzi* for 24 hours. (B) IL-1β levels in the supernatant of bone marrow-derived macrophages primed or not with LPS for 4 hours and further cultured with *T. cruzi* for 24 hours. The cells used in (A) and (B) were collected from wild-type (WT) and *Lgals1^−/−^* mice infected intraperitoneally with *T. cruzi* for 6 hours. Medium: control cells, LPS: lipopolyssacharide from *E. coli*-stimulated cells, Tc: *T. cruzi*. Results are expressed as mean ± standard deviation of three experiments with pooled cells from three mice each. \*\*\**p* < 0.001, \*\**p* < 0.01 and \**p* < 0.05 (unpaired Student’s *t* test).

As macrophages and neutrophils were recruited to the peritoneal cavity of *T. cruzi*-infected WT and *Lgals1^−/−^* mice (Figure 2A), we examined their killing efficiency against the parasites during the early steps of infection. Thioglycolate-elicited macrophages from WT and *Lgals1^−/−^* mice were harvested and cultured with *T. cruzi* isolated from the supernatant of freshly infected LLCMK2 cells. Compared with macrophage cultures *from Lgals1^−/−^* mice, WT mice macrophages released higher number of parasites at four and five days of infection (Figure 6A) and produced less NO at two days of infection (Figure 6B). In the subsequent days, NO levels progressively decreased along time and reached similar levels in macrophage cultures of both WT and *Lgals1^−/−^* mice (Figure 6B).

**Figure 6.**
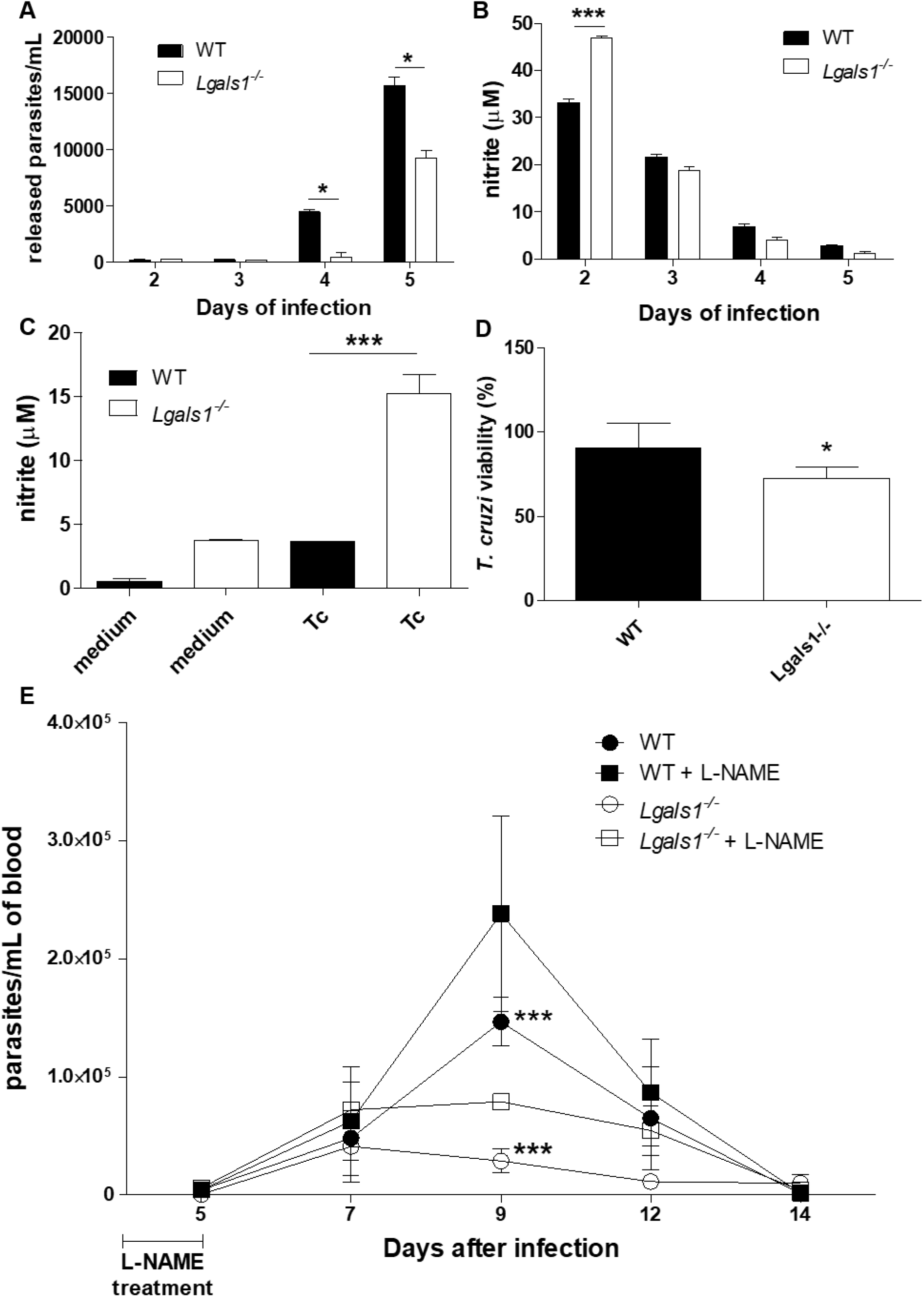
Nitric oxide produced by macrophages and neutrophils from *Lgals1^−/−^* mice controls *T. cruzi* load both *in vitro* and *in vivo*. Thioglycolate-elicited peritoneal macrophages from wild-type (WT) and *Lgals1^−/−^* mice were infected *in vitro* with *T. cruzi* and (A) the number of released parasites and (B) nitrite levels in culture supernatants were determined. Peritoneal cells (>80% neutrophils) were collected from WT and *Lgals1^−/−^* mice infected with *T. cruzi* for 6 hours, and further cultured with *T. cruzi* for 24 hours to measure (C) nitrite levels in the supernatant and (D) the parasite viability. (E) Blood parasitemia after intraperitoneal treatment with L-NAME during the first five days of infection; *** WT *vs Lgals1^−/−^* and *** *Lgals1^−/−^ vs Lgals1^−/−^* + L-NAME, both after 9 days of infection. Results are expressed as mean ± standard deviation (SD) of pooled cells from 3 animals (A, B C and D) and 5 infected animals (E). \**p* < 0.05; \*\**p* < 0.01; \*\*\**p* < 0.001 (unpaired Student’s t test).

To determine NO production by neutrophils, WT and *Lgals1^−/−^* mice were infected with *T. cruzi* and, after six hours, peritoneal cells were collected and cultured with *T. cruzi* for 24 hours *in vitro*. Levels of NO were higher (Figure 6C) and viability of the remaining parasites was lower (Figure 6D) in neutrophil cultures of *Lgals1^−/−^* mice, as compared with neutrophils of WT mice.

Finally, we examined whether treatment with the NO synthase inhibitor L-NAME at early steps (day 0 to 5) of *in vivo T. cruzi* infection affected parasitemia in WT and *Lgals1^−/−^* mice. L-NAME-treated WT and *Lgals1^−/−^* mice exhibited increased blood parasitemia at nine days of infection when compared with their respective non-treated controls (Figure 6E). Blood parasitemia in L-NAME-treated *Lgals1^−/−^* mice was lower than that detected in WT mice, indicating that NO synthase inhibition enhanced parasitemia less effectively than DPI-mediated suppression of ROS production in *Lgals1^−/−^* mice (Figure 4D). Together, ROS and NO production in *Lgals1^−/−^* mice correlated with control of the *T. cruzi* infection at the inoculation site, resulting in decreased parasitemia during the acute phase of this experimental infection.

## 3. DISCUSSION

Our results demonstrate that the lack of Gal-1 in *Lgals1^−/−^* mice is associated with lower parasitemia levels and higher survival rates than their WT counterparts when infected with *T. cruzi* Y strain. We also observed greater neutrophil accumulation in the peritoneal cavity and in heart tissue of *Lgals1^−/−^* mice compared to WT mice. The infiltrated neutrophils in *T. cruzi*-infected *Lgals1^−/−^* mice promoted an increase in the ROS levels, while macrophages and neutrophils produced increased levels of nitric oxide, which limited replication and decreased viability of parasites *in vitro* associated to downregulated IL-1β production. Taken together, our findings demonstrate that a deficiency of Gal-1 suppresses the early steps of *T. cruzi* infection, reducing the pathogenesis and enhancing *T. cruzi*-infected *Lgals1^−/−^* mice survival.

Prior studies have indicated that Gal-1 and Gal-3 may influence *T. cruzi* infection (Benatar et al., 2015; da Silva et al., 2017; Machado et al., 2014; Pineda, Cuervo, Fresno, Soto, & Bonay, 2015; Poncini et al., 2015; Reignault, Barrias, Soares Medeiros, de Souza, & de Carvalho, 2014). As we described here, the lack of Gal-1 promotes a better outcome for the experimental Chagas disease during early infection, corroborating previous a literature report that used a different *T. cruzi* strain (Poncini et al., 2015). In that study, C57BL/6 mice lacking Gal-1 infected with *T. cruzi* strain RA have low mortality rate and muscle parasitemia, probably due to the absence of Gal-1–mediated tolerogenic circuit by dendritic cells, which impairs T cell-dependent anti-parasite immunity (Poncini et al., 2015). The resistance of *Lgals1^−/−^* mice to *T. cruzi* infection reported by that study prompted us to investigate how the lack of Gal-1 affected the early steps of the infection at the peritoneal cavity, focusing mainly on the phagocyte-mediated early responses and neutrophil homeostasis.

Neutrophils are among the immune cells involved in the response against parasitic infections and can remove pathogens by phagocytosis, producing mediators that recruit and activate macrophages and lymphocytes, and releasing molecules that directly control pathogen growth and proliferation, such as reactive oxygen species (ROS), proteases, and neutrophil extracellular traps (NETs) (Scapini and Cassatella, 2014; Sousa et al., 2014; Luna-Gomes et al., 2014; Sousa-Rocha et al., 2015; Dey et al., 2014; Rochael et al., 2015; Koo et al., 2016; Roma et al., 2016). Nevertheless, a tight regulation of inflammatory responses by different mechanisms is essential to protect the host against *T. cruzi* infection without causing tissue damage or other chronic conditions (Chandra et al., 2002; Reyes et al., 2006; Savino et al., 2007).

Considering that the size of parasite inoculum influences the immune response and *T. cruzi* infection outcome in mice (Borges et al., 2013), and that both *Lgals1^−/−^* and WT mice were infected with the same inoculum but had different outcomes, we addressed whether the early response at the infection site influenced the number of parasites that reached the bloodstream and effectively infected the animal tissues. To answer this question, we infected WT and *Lgals1^−/−^* mice intraperitoneally with trypomastigote forms of *T. cruzi*, and analyzed the leukocyte counts in the peritoneal cavity after 6, 24, 48, and 72 hours of infection. Neutrophils were the predominant leukocyte population in the peritoneal cavity of both mice groups at 6 hours of infection, but macrophages predominated at longer infection times (Figure 2A). Compared with WT mice, *Lgals1^−/−^* mice had higher percentage of neutrophils up to 24 hours of infection, and similar leukocyte populations at 48 and 72 hours of infection.

Gal-1 inhibits neutrophil-endothelial cells interaction, decreasing leukocyte influx under inflammatory conditions (La et al., 2003). Neutrophils are crucial to control the first steps of different infections, including those caused by protozoan parasites. For instance, mice infected intradermically with *Leishmania amazonensis* promastigotes exhibit neutrophil accumulation in the lesion between 6 and 24 h of infection, which reduces after 72 hours (Sousa et al., 2014). Neutrophil depletion with specific monoclonal antibodies increases parasite counts and lesion development during the first week of infection with *L. amazonensis* (Sousa et al., 2014). Also, neutrophil depletion one day before infection with *T. cruzi* strain Tulahuén enhances blood parasitemia in BALB/c mice, but reduces blood parasitemia in C57BL/6 mice after 12 and 22 days of infection, respectively; however, it is not clear whether they differ with respect to parasite load at the infection site (Chen, Watanabe, Watanabe, & Sendo, 2001).

Interestingly, as expected, here we demonstrated that the deficiency of endogenous Gal-1 interferes in the early steps of *T. cruzi* infection and favored neutrophil accumulation at the infectious site. Our results indicate that this event promotes a drastic reduction on parasitic load and ameliorates the outcome of experimental Chagas disease, as described under experimental conditions using low parasite inoculum (Borges et al., 2013).

In addition to higher neutrophil counts in the peritoneal cavity in comparison to WT mice, *Lgals1^−/−^* mice heart tissue exhibited elevated neutrophil numbers as indicated by higher MPO activity when compared to WT mice heart tissue after 14 days of infection. In this sense, our findings on neutrophil accumulation into *T. cruzi*-infected mice provide partial explanation for the early control of infection, both at infection site and target organ of the infection. That effect can be explained due to Gal-1 being an endogenous agent for neutrophil homeostasis through its unique property of promoting the phagocytic removal of neutrophil in the absence of apoptosis, a phenomenon called preaparesis (Dias-Baruffi et al., 2003; Stowell, Cho, et al., 2009; Stowell, Karmakar, et al., 2009; Stowell et al., 2007; Stowell et al., 2008).

Our findings corroborate literature reports on the importance of neutrophils to control *T. cruzi* infection. Co-cultures of *T. cruzi*-infected macrophages with neutrophils isolated from C57BL/6 mice display reduced number of trypomastigotes and increased production of TNF-α and NO (Luna-Gomes et al., 2014). Treatment with a selective inducible NO synthase inhibitor favors parasite growth, suggesting that NO plays an important role in parasite killing (Luna-Gomes et al., 2014). Human neutrophils stimulated with live *T. cruzi* or its antigens release NETs, which do not kill the parasites, but immobilize them and interfere with their infection ability (Sousa-Rocha et al., 2015). Mouse neutrophils also produce and release NETs (Etulain et al., 2015), which help to reduce *T. gondii* viability *in vivo* and *in vitro* (Abi Abdallah et al., 2012). Thus, the early and effective neutrophil recruitment to the peritoneal cavity and heart tissue of *Lgals1^−/−^* mice help to control *T. cruzi* infection, but the underlying mechanisms, including NETs release, will be explored in future studies.

Analysis of IFNγ levels in heart tissue from WT and *Lgals1^−/−^* mice after 14 days of infection, when they presented similar inflammatory infiltrates, demonstrated that the cytokine levels were higher in *Lgals1^−/−^* mice (Figure 2C). As IFNγ is an important cytokine to control parasite replication by activating killing mechanisms in both macrophages and neutrophils (Marchi, Sesti-Costa, Ignacchiti, Chedraoui-Silva, & Mantovani, 2014), the strong neutrophil recruitment to the heart of *T. cruzi*-infected *Lgals1^−/−^* mice is linked to the presence of high levels of IFNγ, which improve the control of parasitemia (Figure 1C).

Despite the well-known importance of neutrophils to control parasite infection, especially that caused by protozoans and *T. cruzi* (Abi Abdallah et al., 2012; Luna-Gomes et al., 2014; Sousa-Rocha et al., 2015; Sousa et al., 2014), recruitment of neutrophils and other leukocytes must be tightly controlled to avoid exacerbation of inflammatory responses and tissue damage (Dhiman & Garg, 2011; Paiva, Medei, & Bozza, 2018). In this sense, we examined the profile of leukocyte recruitment to not only the peritoneal cavity but also to heart tissue of *T. cruzi*-infected WT and *Lgals1^−/−^* mice during the acute phase of infection (Figure 3).

It is noteworthy that both WT and *Lgals1^−/−^* mice displayed similar profiles of leukocyte infiltration at 14 days of infection, but they clearly differed at 21 days of infection: the infiltrated cells appeared to be cleared from *Lgals1^−/−^* mice, but they were still present in WT mice. Although the photomicrographs did not distinguish the type of infiltrated cells at 14 days of infection, the increased levels of IFNγ (Figure 2C) and the increased MPO activity (Figure 2 D) in the heart tissue correlated with the lower blood parasitemia in *Lgals1^−/−^* mice (Figure 1C). This tightly regulated recruitment of inflammatory cells to the heart tissue, especially the neutrophil-rich infiltrate in *Lgals1^−/−^* mice, could rapidly clear the parasites that were not killed in the peritoneal cavity and reached the heart. The controlled neutrophil recruitment to the peritoneal cavity and heart tissue, associated with their clearance right after parasite control, prevent exacerbation of inflammatory responses, which could be detrimental to the host.

The fact that *Lgals1^−/−^* mice exhibited higher neutrophil infiltration into the peritoneal cavity and heart tissue, associated with lower parasitemia and longer survival rates, prompted us to investigate the role of ROS production by neutrophils in controling *T. cruzi* replication (Alvarez, Piacenza, Irigoin, Peluffo, & Radi, 2004; Guinazu et al., 2010; Robinson, 2008). Compared with WT mice neutrophils, *Lgals1^−/−^* mice neutrophils produced higher levels of ROS after stimulation with the phorbol ester PMA and the tripeptide fMLP (Figure 4A and 4B, respectively). The NADPH oxidase inhibitor DPI effectively controlled ROS production in fMLP-stimulated *Lgals1^−/−^* mice neutrophils, but failed to control ROS production in PMA-stimulated neutrophils from both WT and *Lgals1^−/−^* mice.

To test the hypothesis that the increased ROS production in *Lgals1^−/−^* mice neutrophils helped to control *T. cruzi* infection at the infection site, we treated WT and *Lgals1^−/−^* mice with DPI solution (1 mg/kg/day, i.p.) during the first five days of infection with *T. cruzi*, and analyzed blood and heart parasite load after the fifth day of infection. At nine days of infection, DPI-treated *Lgals1^−/−^* mice failed to control *T. cruzi* infection, and exhibited circulating parasite levels similar to those detected in WT mice treated or not with DPI (Figure 4C). At 14 days of infection, DPI-treated *Lgals1^−/−^* mice and WT mice treated or not with DPI had high parasite DNA levels in heart tissue (Figure 4D). Only *Lgals1^−/−^* mice not treated with DPI exhibited significantly reduced blood parasitemia at nine days of infection (Figure 4C) and lowered parasite DNA levels at 14 days of infection (Figure 4D).

ROS are key players in the control of parasite infection, especially that caused by *T. cruzi* and *Leishmania* sp. (Roma et al., 2016; Santiago et al., 2012). Here we also examined the role that galectins played in ROS production, but it should be noted that Gal-3 is essential for peritoneal neutrophil ROS production and control of early *T. gondii* infection, and modulates the life span and activation of these leukocytes (Alves et al., 2010; Alves et al., 2013). However, Gal-1, through the neuropilin-1/Smad3 signaling pathway, induces myofibroblast activation, triggers ROS production, and accelerates wound healing (Lin et al., 2015). In our model, recruitment of highly ROS-producing neutrophils to the infection site and heart tissue favors the control of *T. cruzi* proliferation in *Lgals1^−/−^* mice; treatment with the NADPH oxidase inhibitor DPI during early infection *in vivo* mitigated such protective action absence of endogenous Gal-1.

Considering that the increased ROS production by *Lgals1^−/−^* mice neutrophils improved the control of acute *T. cruzi* infection (Figure 4), and that ROS participates in inflammasome activation and IL-1β production (Martinon, 2010; Tschopp & Schroder, 2010), we examined IL-1β production by neutrophils and macrophages from WT and *Lgals1^−/−^* mice, cultured with trypomastigote forms of the parasite. WT mice peritoneal neutrophils released higher IL-1β levels than *Lgals1^−/−^* mice neutrophils (Figure 5A). Similarly, bone marrow-derived macrophages from WT mice, primed or not with LPS and cultured with *T. cruzi*, produced higher levels of IL-1β than *Lgals1^−/−^* mice macrophages (Figure 5B).

Inflammasomes are protein complexes that autocatalytically activate intracellular caspase-1, which cleaves the inactive precursors of IL-1β and IL-18 into bioactive cytokines. IL-1β production is associated with resistance against several infectious agents (van de Veerdonk et al., 2011), including *Leishmania* spp. (Lima-Junior et al., 2013), but it seems not to play an essential role against *T. cruzi*, although some studies have demonstrated its importance to fight against this parasite (Silva et al., 2013). NLRP3 inflammasome controls *T. cruzi* replication in macrophages via caspase-1, independent of IL-1β but dependent on NO production (Goncalves et al., 2013). Macrophages from *T. cruzi*-infected *NLRP3^−/−^* mice produce high basal levels of ROS, which contribute to parasite killing through an IL-1β-independent pathway (Dey et al., 2014).

Several of our results were unexpected, including the observation that WT neutrophils and macrophages release higher levels of IL-1β than *Lgals1^−/−^* leukocytes when in contact with *T. cruzi* (Figure 5). By contrast, WT leukocytes produced lower levels of ROS and NO than *Lgals1^−/−^* leukocytes (Figures 4 and 6). Production of oxidant species is the first line of defense against *T. cruzi* before IL-1β production or action (Dey et al., 2014; Goncalves et al., 2013). Hence, *Lgals1^−/−^* resistance to *T. cruzi* can be directly related to a rapid and regulated ROS and NO production.

Several studies have reported that NO controls replication of intracellular parasites, especially *T. cruzi* (Aliberti, Machado, Gazzinelli, Teixeira, & Silva, 1999; Costa et al., 2006; Vespa, Cunha, & Silva, 1994). Here we measured the levels of NO production in WT and *Lgals1^−/−^* phagocytes stimulated with *T. cruzi* (Figure 6). *Lgals1^−/−^* macrophages and neutrophils were more efficient in producing NO and killing *T. cruzi* than the WT leukocytes (Figures 5A and 5B). To examine the role that NO production played *in vivo* during early *T. cruzi* infection, *Lgals1^−/−^* and WT mice were treated with the NO synthase inhibitor L-NAME before infection. At the ninth day of infection, L-NAME-treated *Lgals1^−/−^* and WT mice exhibited increased blood parasitemia when compared with their respective untreated controls (Figure 6E). Blood parasitemia in L-NAME-treated *Lgals1^−/−^* mice was lower than that detected in both groups of WT mice, probably due to the enhanced ROS production in *Lgals1^−/−^* mice during early infection.

NO production can be detrimental to the mammalian host during *T. cruzi* infection, especially at chronic phase (Carvalho et al., 2012; Tatakihara et al., 2015). In addition, the lack of Gal-1 is related to stronger NO production, neutrophil infiltration, and release of inflammatory cytokines in mice infected with the fungus *Histoplasma capsulatum*, resulting in lower survival rates (Rodrigues et al., 2016). Alternatively, NO helps to regulate ROS production and protects against septic shock (Mao et al., 2013). NO inhibits the NLRP3-mediated ASC pyroptosome formation, caspase-1 activation, and IL-1β secretion in myeloid cells from both mice and humans. Depletion of inducible NO synthase stimulates IL-1β production and caspase-1 activation, which enhances NLRP3-dependent cytokine production *in vivo* and thereby increases mortality from LPS-induced sepsis in mice (Mao et al., 2013). Therefore, excessive IL-1β production not associated with tight control of NO production can culminate in death of infected mice due to exacerbation of inflammatory process. In the present study, WT leukocytes produced more IL-1β (Figure 5) and less ROS and NO (Figures 4 and 6, respectively) than *Lgals1^−/−^* leukocytes. In addition, WT mice exhibited higher mortality rate than *Lgals1^−/−^* mice (Figure 1), which is probably due to a regulated NO production early during infection. Taken together, the findings reported herein suggest that rapid neutrophil recruitment associated with increased NO and ROS release protect *Lgals1^−/−^* mice during *T. cruzi* acute infection.

## 4. EXPERIMENTAL PROCEDURES

### 4.1 Mice and ethics

Wild type (WT) C57BL/6 mice were purchased from the Central Animal Facility at Ribeirão Preto campus of the University of São Paulo, Ribeirão Preto, SP, Brazil. C57BL/6 mice lacking Gal-1 gene (*Lgals1^−/−^*) were obtained from the Animal Facility at School of Pharmaceutical Sciences of Ribeirão Preto, University of São Paulo, Ribeirão Preto, SP, Brazil. WT and *Lgals1^−/−^* mice weighed 20-25 g at the moment of the experiments. The Ethics Commission on Animal Use (CEUA) at Ribeirão Preto campus of the University of São Paulo, Ribeirão Preto, SP, Brazil approved the study protocol (n° 08.1.587.53.1).

### 4.2 Parasites maintenance and isolation

Trypomastigote forms of *T. cruzi* strain Y were maintained by serial weekly passages in Swiss mice. Blood was collected in the acute phase of infection (6 to 8 days after infection), and about 10^6^ trypomastigotes were freshly separated and used to infect sub-confluent LLC-MK2 cells (American Type Culture Collection, ATCC, CCL7.1) maintained in 75 cm^2^ bottles, at 37°C and 5% CO_2_ in RPMI1640 medium containing 10% fetal bovine serum (FBS), 1% glutamine, 100 UI penicillin, 100 μg of streptomycin, and 0.25 μg/mL of amphotericin B per mL (complete medium - all reagents from Sigma-Aldrich). To separate the trypomastigote forms from the culture, the supernatant was centrifuged at 1,000 ×*g* for 30 minutes, at 4°C.

### 4.3 Mice infection, parasitemia, and survival

C57BL/6 WT and *Lgals1^−/−^* mice (six animals per group) were infected intraperitoneally with 1,000 *T. cruzi* trypomastigote forms freshly separated from blood of acutely infected Swiss mice, and diluted in 200 μL of sterile phosphate-buffered solution (PBS). Blood parasite counts were determined as described by (Brener, 1962). Briefly, 5 μL of freshly collected tail blood were placed in a slide and covered with a coverslip to count parasites in 50 microscopic fields (400× magnification), every 2-3 days from 5 to 14 days after infection. To determine the survival rate, both infected mice groups were monitored daily until 60 days of infection.

### 4.4 T. cruzi DNA quantification in heart tissue by real-time PCR

Approximately 30 mg of heart tissue from WT and *Lgals1^−/−^* mice were washed with PBS. The genomic DNA was isolated using *Illustra™ Tissue and cell genomic Prep mini spin kit* (GE, USA), according to the manufacturer’s instructions. The amount of recovered genomic DNA was quantified with *NanoDrop ND 2000* (Thermo Scientific, USA) and stored at −20 °C. Amplification was performed using 5 μL of specific *SybrGreen Master Mix reagent* (Thermo Scientific), 0.35 μM of each specific primer for *T. cruzi* genomic DNA (TCZ-F: GCTCTTGCCCACAAGGGTGC 5’-3’ and 5’-TCZ R CCAAGCAGCGGATAGTTCAGG-3’) (Caldas et al., 2012; Cummings & Tarleton, 2003), 10 ng of DNA from each heart sample, and nuclease-free water sufficient to make up the reaction volume to 10 μL. A *StepOne ™ Real Time PCR System* (Life Technologies, USA) apparatus was used to obtain C_T_ values. DNA samples from non-infected mice were used as the negative control. A standard curve with dilution series of genomic DNA obtained from culture-derived *T. cruzi* was used to calculate the relative number of parasites in 10 ng of sample DNA.

### 4.5 T. cruzi-induced peritonitis

The morphology of immune cells that migrated to the peritoneum after 6, 24, 48, and 72 hours of infection of WT and *Lgals1^−/−^* mice with 1,000 trypomastigote forms of *T. cruzi* was analyzed as follows. Further to euthanasia peritoneal cells were collected by injecting and aspirating 5 mL of ice-cold sterile PBS into and from the cavity, using syringe and needle. Cells were centrifuged at 300 ×*g* for 10 minutes at 4°C, and the pellet was suspended in cold PBS. Fifty microliters of this suspension were placed on a microscope slide, centrifuged (Shandon Cytospin), dried, fixed, and stained with *Panótico Rápido* (Laborclin, Brazil). The percentage of each cell type was estimated by optical microscopy.

### 4.6 Myeloperoxidase (MPO) and N-acetyl-β-D-glucosaminidase (NAG) activity in cardiac tissue

The measurement of MPO and NAG activity, enzymes predominantly found in neutrophils and macrophages, respectively (Ayala et al., 2000), was employed to detect these cells in heart tissue. Briefly, heart tissue was homogenized in cold PBS (Ultra-Turrax disperser – IKA), centrifuged, and red blood cells in the tissue pellet were lysed by osmotic difference adding hypotonic NaCl solution. The homogenate was centrifuged again, and the pellet was suspended in PBS with 0.5% (w/v) of hexadecyltrimethylammonium bromide (Sigma-Aldrich) under vigorous stirring, frozen and thawed three times, and centrifuged at 10,000 ×*g* for 15 minutes at 4°C. The resulting supernatant was stored at −80°C until used. To determine MPO and NAG activity. To determine MPO activity, 50 μL of supernatant was mixed with 50 μL of the MPO substrate 3,3’,5,5’-tetramethylbenzidine (Sigma-Aldrich) in a 96-well microplate. After a 30-minute incubation at 37°C, in the dark, the reaction was stopped by adding 25 μL of a 4 M H_2_SO_4_ solution to each well, and the absorbance was recorded at 450 nm. To determine NAG activity, 25 μL of supernatant was mixed with 25 μL of *p*-nitrophenyl-2-acetamide-β-D-glucopyranoside and 100 μL of citrate buffer in a 96-well microplate. After a one-hour incubation at 37°C, in the dark, the reaction was stopped by adding 100 μL of glycine buffer to each well, and the absorbance was recorded at 405 nm. The absorbance values at 450 nm and 405 nm were proportional to the tissue MPO and NAG activity, respectively. The activity of each enzyme per gram of tissue was obtained by dividing the absorbance values (after discounting the blank) by the total tissue mass that was initially homogenized.

### 4.7 Inflammatory peritoneal neutrophils from T. cruzi-infected mice

After 6 hours of intraperitoneal *T. cruzi* infection, neutrophils were obtained by gently injecting and aspirating 5 mL of cold sterile PBS into and from the peritoneal cavity of mice. The suspension was centrifuged at 300×*g* for 10 minutes, at 4°C, and the neutrophil-rich pellet was treated with erythrocyte lysis buffer (0.16 M NH_4_Cl and 0.17 M Tris-HCl, pH 7.5) for 5 minutes at 37°C, when appropriate. After washing with ice cold PBS, cells were suspended in complete RPMI medium and viable cells were counted in a hemocytometer by trypan blue exclusion. Neutrophils obtained in this manner were routinely >95% viable and 80–90% pure.

### 4.8 Reactive oxygen species (ROS) production

Chemiluminescence assay was performed according to the protocol reported by (Alves, Marzocchi-Machado, Carvalho, & Lucisano Valim, 2003; Cheung, Archibald, & Robinson, 1983). Inflammatory peritoneal neutrophils (5×10^5^ cells/500 μL) obtained from *T. cruzi*-infected mice after 6 hours of infection were incubated with 5 μL of 280 μM luminol in Hanks’ balanced saline solution (HBSS) for 2 min at 37°C. Cells were then stimulated with 50 μL of 10^−7^ M phorbol 12-myristate 13-acetate (PMA, Sigma-Aldrich) or N-formyl-methionyl-leucyl-phenylalanine (fMLP, Sigma-Aldrich), and chemiluminescence was recorded for 20 or 2 minutes, respectively, at 37°C, in a luminometer (Auto Lumat LB 953 EG & G Berthold, Bad Wildbad, Baden-Württemberg, Germany). Non-stimulated cells incubated with luminol were used as controls for spontaneous (basal) ROS release. Diphenyleneiodonium chloride (DPI, 10 μM), a NADPH oxidase inhibitor, was used as the negative control for ROS release *in vitro* (Bansal et al., 2012). The spontaneous ROS release was subtracted from the samples and the results were expressed as the integrated area of chemiluminescence profiles (area under the curve).

### 4.9 T. cruzi viability in inflammatory peritoneal neutrophils cultures

To assess IL-1β and NO production, inflammatory peritoneal neutrophils obtained from *T. cruzi*-infected WT and *Lgals1^−/−^* mice, after 6 hours of infection, were washed with HBSS, suspended in complete RPMI medium (5×10^5^ cells/500 μL), and co-cultured (24 hours, 37°C, humidified atmosphere with 5% CO_2_) with trypomastigote forms of *T. cruzi* freshly isolated from infected LLCMK2 cell culture supernatant (5 parasites: 1 neutrophil). The well content was removed and centrifuged at 300 ×*g* for 5 minutes (to separate remaining neutrophils), and the resulting supernatant was recovered and centrifuged in a new tube at 1,000 ×g for 30 minutes at 4°C (to recover the parasites). Then, the supernatant was stored at −80 °C until determination of IL-1β and nitrite concentrations, while the *T. cruzi*-rich pellet was suspended in 100 μL of fresh complete medium and incubated for 24 hours at 37°C in a humidified atmosphere with 5% CO_2_, in a 96-well culture plate. To determine parasite viability, 5 μL of a 10 mg/mL solution of resazurin sodium salt (Sigma-Aldrich) prepared in PBS (pH 7) were added to each well and the plate was incubated for 4 hours at 37°C (Rolon, Vega, Escario, & Gomez-Barrio, 2006). The absorbance was recorded at 570 and 600 nm in a SpectraMax Microplate Reader (Molecular Devices). Control wells with parasites cultured in the absence of neutrophils represent 100% *T. cruzi* viability.

### 4.10 In vivo treatment with DPI and L-NAME

In some experiments, WT and *Lgals1^−/−^* mice were treated intraperitoneally with 200 μL of DPI (1 mg/kg/day) or Nω-Nitro-L-arginine methyl ester hydrochloride (L-NAME, 100 mg/kg/day) from day 0 to day 5 after infection with *T. cruzi*. Solutions containing the necessary amount of each inhibitor were freshly prepared daily in sterile PBS.

### 4.11 Bone marrow-derived macrophages (BMDM) and IL-1β measurement

BMDM were obtained from WT and *Lgals1^−/−^* mice as described by(Marim, Silveira, Lima, & Zamboni, 2010). Briefly, bone marrow from both femurs of three animals were removed with RPMI medium supplemented with 10 mM of L-glutamine, 30% LCCM medium (*L929-cell conditioned medium*) containing macrophage colony stimulating-factor, and 10% FBS. Cells were cultured in BD Optilux polyestirene Petri dishes (4×10^6^ cells/mL) for 7 days at 37 °C and 5% CO_2_, with subsequent medium addition after 4 days. The macrophages were then detached from the dishes, washed with sterile PBS and cultured in a 24-well plate (5×10^5^ cells/500 μL) for 24 hours. Next, the cells were primed for 4 h with 500 ng/mL ultrapure LPS (InvivoGen). The supernatant was discharged and macrophages were infected with trypomastigote forms of *T. cruzi* freshly isolated from infected LLCMK2 cell culture supernatant (5 parasites: 1 neutrophil). After incubation (24 hours, 37°C, humidified atmosphere with 5% CO_2_), the cell suspensions were centrifuged (300 ×*g*, 10 min, 4 °C) and the resulting supernatants were collected and stored at −80°C. The concentration of IL-1β was determined using an ELISA kit (BD Biosciences).

### 4.12 Trypanocidal activity in infected peritoneal macrophages

WT and *Lgals1^−/−^* mice were treated with intraperitoneal injection of 1 mL of sterile 3% thioglycolate solution and, after four days, mice were euthanized and 5 mL of cold PBS were injected into and aspirated from the peritoneal cavity of the animals. The exudate was washed twice (300 ×*g*, 5 minutes, 4°C), and the resulting cell pellet was suspended in complete medium. The cells were counted, distributed into a 24-well plate (5×10^5^/500 μL/well), incubated for 24 hours at 37°C in a humidified atmosphere with 5% CO_2_, and washed three times with RPMI medium to remove non-adherent cells. Trypomastigote forms of *T. cruzi* obtained from infected LLCMK2 cell culture supernatant (5 parasites: 1 macrophage) were suspended in 500 μL RPMI and added to the wells (Talvani et al., 2002). After 3 hours of incubation, the non-internalized parasites were removed by successive washes with RPMI medium. The plates were incubated for five days under the abovementioned conditions, with daily collection of culture supernatants and addition of fresh medium to the wells. Such supernatants were centrifuged (1,000 ×*g*, 4°C, 10 minutes), and the resulting pellet was suspended in 50 μL of a 0.1% trypan blue/10% formaldehyde (v/v) solution to count the released trypomastigotes in a Neubauer chamber.

### 4.13 NO determination

To determine NO levels in the supernatant of different cultures, the concentration of nitrite was determined by the Griess reaction (Gilliam, Sherman, Griscavage, & Ignarro, 1993; Vespa et al., 1994). Briefly, 50 μL of each culture supernatant were mixed with 50 μL of Griess reagent [1:1 (v/v) 0,1% *N*-(1-naphthyl) ethylenediamine dihydrochloride (NEED, Sigma-Aldrich) in water: 1% sulfanilamide (Sigma-Aldrich) in 3% phosphoric acid (Sigma-Aldrich)] in 96-well plates (Costar). After 5 minutes of incubation, the absorbance was recorded at 554 nm in a spectrophotometer. RPMI was used as blank reaction and a calibration curve was made with known concentrations of NaNO_2_ (Sigma-Aldrich) diluted in RPMI. The assay was performed in triplicate.

### 4.14 Statistical analysis

Experimental data were processed and analyzed using the GraphPad Prism software, version 5 for Windows (GraphPad Software Inc., USA). Two-way analysis of variance (ANOVA) combined with Bonferroni *post-hoc* test was used to compare three or more experimental groups, while unpaired Student *t* test was used to compare two groups at a single point. *p*<0.05 was considered significant.

## ACKNOWLEDGEMENTS

This work was supported by The São Paulo Research Foundation (FAPESP, grant n° 2010/10470-6) and Brazilian Council for Scientific and Technological Development (CNPq – 312606/2019-2 and 432201/2016-5). This study was partially funded by Coordenação de Aperfeiçoamento de Pessoal de Nível Superior – Brasil (CAPES) – Código de Financiamento 001.

